# Iterative prioritisation of upstream and downstream passage remediation for the critically endangered European eel (*Anguilla anguilla*) at pumping stations

**DOI:** 10.64898/2026.11.15.766773

**Authors:** Angus I.T. Monaghan, Nathan P. Griffiths, Andy D. Nunn, Graham S. Sellers, Rosalind M. Wright, Jonathan D. Bolland

## Abstract

The European eel (*Anguilla anguilla*) is critically endangered and subject to regulations requiring safe passage at migratory barriers and water intakes, but remediation must be prioritised to maximise conservation benefits. This study developed a novel prioritisation framework for both upstream and downstream eel passage remediation in heavily modified and fragmented pumped river systems. Catchment area (habitat quantity), distance from the tidal limit and numbers of migration barriers downstream were used as baseline geospatial metrics for bi-directional passage data. Following this, eel presence/absence data was incorporated to separate sites into upstream and downstream passage prioritisations, with site-specific hydrological connectivity (downstream) and fish community richness (upstream) data further refining rankings. Prioritisation rankings were compared iteratively to assess the influence of combining variables. Eel presence/absence data substantially altered downstream passage priorities relative to geospatial metrics alone, with a maximum rank change of 96 and eleven sites changing by 50 or more places. For downstream passage, hydrological connectivity resulted in significant re-ranking of priorities. For upstream passage, including fish species richness as an indicator for habitat quality caused notable re-ranking of sites compared with geospatial metrics alone. The results demonstrate that prioritisation based solely on geospatial metrics risks overlooking important sites and, consequently, directing resources towards areas with limited conservation potential. The iterative, flexible framework presented in this study provides a transparent and transferable approach for informing eel passage investment within heavily modified drainage systems and could be adapted to other landscapes where anguillid eels are conservation priorities.

## Introduction

The European eel (*Anguilla anguilla*, hereafter referred to as “eel”) is a catadromous fish species found across Europe. To complete their life cycle, individuals colonise freshwater, estuarine or coastal habitats as juveniles and migrate to the spawning grounds in the Sargasso Sea as adults. The species has experienced an approximately 95% decline in recruitment in the last 50 years and, consequently, is classified as critically endangered with extinction (Pike et al., 2020). This decline is the result of a multitude of factors, including overfishing, habitat fragmentation and degradation, climate change, pollution and hindrance to spawning migrations (Dekker, 2000; Pike et al., 2020). To aid their protection, the EU introduced legislation for conservation measures with the development of Eel Management Plans (EC Regulation No. 1100/2007; European Commission, 2007). The Eels (England and Wales) Regulations 2009 has specific legislation that requires any intakes abstracting >20 m^3^ of water per day, including pumping stations, to install European eel protection measures. Non-compliant pumping stations currently have exemptions issued by the regulator (i.e. the Environment Agency) until mitigation can be achieved during a non-eel-related programme of works to replace existing pumps.

Eels are known to be abundant in low-lying wetlands near the coast (Laffaille et al., 2004), many of which have been reclaimed in recent centuries by land drainage measures, which often make use of pumping stations. Drainage and subsequent anthropogenic land use transforms low-lying landscapes from dynamic marshlands to homogenous matrixes of agricultural fields and strictly defined drainage channels (Van Loon et al., 2009). Agricultural runoff, a lack of flow during summer months and intensive riparian and in-channel management (Swales, 1982; Montgomery et al., 2018) leave drainage systems vulnerable to thermal and water quality-induced mortality events (Lu et al., 2023). In turn, the resulting floral and faunal communities differ from those that would occur naturally (Townsend & Peirson, 1988; Blann et al., 2009). Pumping stations pose a particular threat to eels as they are a barrier to upstream habitat colonisation by juveniles (Griffiths et al., 2025a) and downstream (seaward) spawning migrations of mature “silver” eels (Bolland et al., 2019). Furthermore, eels and other fish can be injured and killed when entrained in stations with traditional pump mechanisms (Perry et al., 2015; Norman et al., 2023a, 2023b).

To overcome the negative effects of pumping stations on eels, upstream migration can be mitigated with elver passes, gravity sluices or trap and transport (Solomon et al., 2004), and downstream migration can be mitigated with gravity sluices and pump mechanisms intended to be fish-friendly (also referred to as “less damaging” (Carter et al., 2023b; Evans et al., 2024). However, it is not logistically or fiscally feasible to modify all pumping stations in a short period of time. Furthermore, in the absence of present-day knowledge of eel occurrence in understudied pumped catchments, it must be assumed eels are present upstream of all pumping stations given their historic distribution (Porter, 1958). Consequently, in England and Wales, when upstream and downstream eel passage mitigation is implemented during routine work to replace existing pumps (to ensure regulatory compliance), an unknown fraction have no eels in the upstream catchment. Even in the absence of eels, fish-friendly pumps are still preferred over traditional alternatives given the potential benefits to the wider fish community, although it must be acknowledged that eel-friendly pumps may not be safe for other all fish species (Bruneel et al., 2024). Furthermore, there is a risk that upstream passage may be provided into catchments with degraded upstream habitats and thus may unintentionally lead to eel mortalities. Conversely, there may be catchments with large numbers of eels and/or high-quality habitat upstream of pumping stations that are not due for refurbishment for decades. Consequently, efforts are urgently required to perform targeted and optimised eel protection across pumping stations at a large scale.

Across the world, freshwater ecosystems are highly fragmented by artificial barriers to fish movement (Belletti et al., 2020; Parasiewicz et al., 2023). In response, regulatory authorities, conservation organisations and infrastructure owners are faced with the challenge of strategically remediating connectivity to achieve the greatest ecological benefit, invariably with limited resources. To avoid *ad hoc* or opportunistic site-specific interventions informed only by local knowledge that fail to maximise conservation benefit, prioritisations can be performed to score and rank multiple barriers across broad spatial scales. However, approaches are highly variable depending on target species, infrastructure type and data availability, which influence strengths, limitations and transferability. In addition, the majority of peer-reviewed passage prioritisation frameworks have considered only immigrating fishes at low-head barriers (Kocovsky et al., 2009; Kemp & O’hanley, 2010; Nunn & Cowx, 2012), with pumping stations and emigrations receiving little attention.

Despite their challenging catadromous life-history strategy and many species being of conservation concern (Pike et al., 2020), no peer-reviewed passage prioritisations appear to have focused exclusively on anguillid eels or considered immigrating and emigrating life-stages separately to tailoring conservation measures. Indeed, to our knowledge, Kocovsky et al. (2009) and Nunn & Cowx (2012) are the only peer-reviewed prioritisations to have included anguillid eels, merely considering upstream passage among all prevailing diadromous species. Furthermore, no peer-reviewed prioritisations appear to exist for pumping stations, despite their considerable hydroecological impacts coupled with the significant cost and engineering challenge associated with retrofitting fish protection at existing sites. An unpublished regional-scale pumping station (n = 444) prioritisation was performed by Solomon & Wright (2012) using only geospatial metrics (i.e. distance from tidal limit and catchment area), again predicated on the assumption that European eels were historically present upstream of all pumping stations. Consequently, some sites may have been prioritised for passage improvements despite supporting no eels and/or poor-quality habitat, while sites with greater conservation potential may have been overlooked. To address this possibility, the fish community upstream of a sub-set of pumping stations in the region (n = 152) has since been assessed using environmental DNA metabarcoding; European eels were present upstream of less than 30% of pumping stations (Monaghan et al., 2026). Abiotic and biotic drivers of eel occurrence in pumped river catchments were also revealed, which have implications for prioritising infrastructure management for European eel conservation.

Fish passage prioritisation studies commonly justify, define and weight variables and report the resulting rankings (Kocovsky et al., 2009; Kemp & O’hanley, 2010; Nunn & Cowx, 2012). Indeed, to our knowledge, no previous fish-focused studies have explicitly quantified how priorities change as progressively more detailed sources of information are incorporated, as has been performed for terrestrial conservation prioritisations (Kujala et al., 2018). This may be attributed to fish prioritisations typically being desk-based studies using geospatial data, rather than incorporating fish community data or site-specific information (but see Nunn & Cowx, 2012; King & O’Hanley, 2016). In the present study, prioritisation was developed iteratively, beginning with geospatial metrics, i.e. distance from the tidal limit and catchment area, and the numbers of downstream barriers (Stage 1). Comprehensive, targeted and contemporary eel occurrence data were then incorporated to uniquely prioritise upstream and downstream eel passage separately (Stage 2). Thereafter, the number of downstream barriers (derived from satellite imagery), site-specific operational data and fish community composition (as a proxy for habitat quality) were integrated to further refine the prioritisations (Stage 3). The iterative process allowed the influence of each additional source of information on site priorities to be quantified and visualised, rather than assessing only the final composite prioritisation.

The overall aim of this study was to uniquely prioritise upstream and downstream eel passage remediation at pumping stations at ecosystem scale. The study objectives were to assess:

- the interrelationship between geospatial metrics on bi-directional passage prioritisation at all pumping stations (Stage 1),
- the influence of eel-presence to inform targeted directional (upstream and downstream) eel passage remediation (Stage 2),
- the influence of site-specific hydrological connectivity and wider fish community on downstream and upstream prioritisations, respectively (Stage 3), and
- the influence of iteratively incorporating additional sources of information on the final prioritisations.

The outputs of this study will simultaneously inform landscape-scale eel conservation and pumping station management while also providing a reproducible prioritisation framework that could be applied globally to low lying areas with anguillid species.

## Methods

### Study region

This study focused on 149 pumping stations in The Fens, East Anglia, England; a 400,000 hectare marshland in the lower reaches of the rivers Great Ouse and Nene catchments, which has been drained since the 17^th^ century for agricultural and flood defence purposes (Williams & Worth, 2003). The objectives of upstream and downstream eel passage prioritisation in The Fens differ and therefore require different approaches. Downstream passage prioritisation aims to maximise the safe emigration of adult eels from where they are most abundant. Conversely, upstream passage prioritisation aims to maximise the immigration of juvenile eels to currently inaccessible areas where habitat quality is highest. As direct estimates of habitat quantity and quality, as well as upstream and downstream eel abundance, were unavailable, a series of geospatial, biological and site-specific variables were used as proxies (Table 1) in three stages. Geospatial variables were included in both upstream and downstream prioritisation frameworks (Stage 1), with specific variables used to refine directional passage priorities according to their ecological relevance (Stage 3) following incorporation of present-day eel distribution (Stage 2).

**Table 1.** Factors considered for prioritising eel passage, their associated code, migration direction (Bi = bi-directional, Down = downstream and Up = upstream), rationale and score preparation, including formula used (V = value and MV = maximum value).

| Factor | Code | Direction | Proxy and rationale | Score preparation |
|---|---|---|---|---|
| Catchment area | CA | Bi | A proxy for habitat quantity; larger catchments provide more habitat and therefore have the potential to support larger eel stocks | Natural log transformed and scaled from 1-5; largest catchment scored 5. Formula: 6-(((V/MV)*4)+1) |
| Distance from tidal limit | DTL | Bi | A proxy for downstream eel abundance; probability of eels being present at a pumping station decreases with increasing distance from the tidal limit | Inverted and scaled from 1-5; pumping stations at the tidal limit scored 5. Formula: ((V/MV)*4)+1) |
| Barriers Downstream | BD | Bi | A proxy for downstream eel abundance; probability of eels being present at a pumping station decreases with an increasing number of barriers between tidal water | Scored as a penalising multiplier reflecting the increasing effect of multiple barriers:<br>• 0 barriers = 1<br>• 1 barrier = 0.9<br>• 2 barriers = 0.75<br>• 3 barriers = 0.5 |
| Eel presence | EP | Bi | Evidence of presence; downstream passage remediation should be prioritised at pumping stations with eels and station without eels should be prioritised for upstream passage remediation | Sites without eels were excluded from downstream passage prioritisation and instead prioritised for upstream passage remediation. |
| Hydrological connectivity type | HCT | Down | A proxy for upstream eel abundance; greater connectivity between a pumping station and the downstream river network is likely to result in greater numbers of eels colonising habitat upstream of the station. | Connectivity scores from Monaghan et al. (2026) spaced evenly from 1-5 |
| Winter fish species richness | WFR | Up | A proxy for upstream habitat quality; biotic indicator of physical habitat and water quality | Scaled from 0-1; highest richness scored 1. Formula: V/MV |

### Stage 1. Geospatial bi-directional passage prioritisation

Sites were initially ranked according to only geospatial metrics to understand priorities without knowing eel distribution. Consequently, simultaneous remediation of bi-directional passage is inherently recommended.

#### Catchment area (CA); a proxy for habitat quantity

It was assumed that the ecological benefits of passage improvements would be maximised by facilitating immigration into and emigration from areas with the potential to support the largest numbers of eels. Thus, catchment area (CA) was used as a proxy for habitat quantity. Monaghan et al. (2026) also revealed eel presence was positively associated with catchment area. CA was natural log transformed (due to values spanning several orders of magnitude) and scaled from 1-5, with large catchments scoring highest.

#### Distance from tidal limit (DTL); a proxy for downstream eel abundance

The probability of catadromous species reaching an inland location is intrinsically and negatively linked to their distance from the tidal limit (DTL) (Ibbotson et al., 2002; Griffiths et al., 2023; Monaghan et al., 2026). Thus, DTL was used as a proxy for downstream eel abundance. It is also likely that the abundance upstream of a pumping station would be greater the closer it is to tidal waters and thus a higher priority for safe downstream passage remediation than stations further inland. Indeed, it was assumed the ecological benefits of upstream and downstream passage improvements will be maximised the closer a pumping station is to tidal water. DTL was inversely scaled from 1-5, with sites at the tidal limit scoring highest.

#### Barriers downstream (BD); a proxy for downstream eel abundance

Migration barriers negatively affect eel distribution and abundance (Monaghan et al., 2026), with each successive barrier having a disproportionately greater negative effect on immigration (Yokouchi et al., 2022). Thus, the number of barriers downstream (BD), i.e. between the pumping station and tidal water, was used as a proxy for downstream eel abundance. It is also likely that barriers will impede seaward migration (Verhelst et al., 2025) and are therefore considered lower priority for downstream passage. Indeed, it was assumed that the ecological benefits of passage improvements will be maximised by facilitating immigration into and emigration from pumping stations with the fewest barriers downstream. Sites with zero (pumping stations discharging into tidal water), one, two and three barriers downstream were allocated multiplier scores of 1, 0.9, 0.75 and 0.5, respectively. The passability of migration barriers were not empirically derived. Instead, the scoring reflects the increasingly negative effect of successive barriers (Yokouchi et al., 2022) and thus down weights sites with multiple barriers downstream.

### Stage 2: Eel presence-informed directional passage prioritisation

Comprehensive, targeted and contemporary eel occurrence data (Monaghan et al., 2026) were incorporated to prioritise upstream and downstream eel passage separately. Pumping stations where eels were detected upstream were prioritised for downstream passage to protect existing eel stocks. Sites where eels were not detected upstream of the pumping station were considered for upstream passage prioritisation to improve connectivity and increase habitat availability for immigrating juvenile eels.

### Stage 3. Prioritisation refinement using site-specific features

#### Downstream passage. Hydrological connectivity type (HCT); a proxy for upstream eel abundance

It is desirable to prioritise downstream passage at pumping stations with the largest numbers of eels upstream. However, it is not possible to estimate eel abundance from eDNA data (Rourke et al., 2022). Monaghan et al. (2026) revealed that eel occurrence was positively associated hydrological connectivity type (HCT), and thus it was used as a proxy for eel abundance, i.e. sites with better upstream connectivity are likely to contain more eels. Scores were scaled evenly between 1-5; sites with no connectivity were assigned a score of 1, sites with manual connectivity (i.e. gravity-fed irrigation inlets) were assigned a score of 2.33, sites prone to flooding from the downstream waterbody were assigned a score of 3.66 and sites with gravity drainage in place were assigned a score of 5.

#### Upstream passage. Winter fish richness (WFR); proxy for upstream habitat quality

Despite eels’ generalist and tolerant nature compared to many fish species (Nälsund et al., 2022), facilitating upstream passage into new habitats, where individuals may reside as yellow eels for many years, should prioritise those of the highest quality and greatest resilience (Nunn & Cowx, 2012). This is particularly relevant as the continued effects of climate change, such as rising temperatures, may further degrade drain habitats in the future (Rideout et al., 2022). Fish communities can be used as reliable, long-term biotic indicators of habitat quality (Pont et al., 2006; Noble et al., 2007). Consequently, fish community data from eDNA surveys (Monaghan et al., 2026) were used in this study as a proxy for upstream habitat quality. Specifically, due to the potential for summer water transfers (e.g. for irrigation) to move fish and/or fish DNA that do not persist in poor quality pumped catchments, median sample richness from eDNA samples collected in March (n = 75 sites) or 1-sample fish species richness (n = 31 sites) were used to derive a winter fish species richness (WFR). WFR was scaled from 0-1, with sites with richer fish communities scoring highest.

### Data analysis

All continuous variables that were scaled either 0-1 or 1-5 were not rounded and raw scaling scores were retained. Prioritisation scores were calculated multiplicatively and converted to ordinal ranks, with rank 1 representing the highest priority site. Passage prioritisation scores were developed iteratively to quantify the effect of each new variable on the overall site rankings. Spearman’s rank tests were used to test the strength, direction and significance of correlations between successive iterations. Kruskal-Wallis tests with post-hoc Dunn tests and Bonferroni corrections were used to test for differences between DTL and BD, to determine whether these variables were strongly associated and therefore potentially represented overlapping measures of connectivity. Mapping of sites by passage priority was carried out in a custom workflow in R. All data analysis was carried out in R version 4.5.1 (R Core Team, 2025) and utilised the following packages: Tidyverse (Wickham et al., 2019) and dunn.test (Dinno, 2014). Map visualisation used the custom R-based workflow of Monaghan et al. (2026).

## Results

### Stage 1. Geospatial bi-directional passage prioritisation

Sites ranked for bi-directional passage based upon CA or DTL alone exhibited low levels of agreement. There was a weak, non-significant correlation between the ranks (Spearman’s rank, rho = 0.074, p = 0.37) (Figure 1). When combined, the “CAxDTL” score showed a stronger correlation with the DTL alone ranking (Spearman’s rank, rho = 0.94, p < 0.001) than with CA alone (Spearman’s rank, rho = 0.35, p < 0.001), due to the higher variance in DTL than CA and, thus, this metric driving the score more than CA (Figure 1). When BD was introduced to the formula (CAxDTLxBD) for bi-directional passage prioritisation, agreement remained robust, with barriers having an approximately 1% effect on rank change (Spearman’s rank, rho = 0.99, p < 0.001) and re-ranking was minimal (largest rank shift was 23, Figure 1). DTL increased significantly with downstream barrier number (Kruskal-Wallis test, X² = 43.20, df = 3, p < 0.001), showing the two factors are closely linked.

**Figure 1.**
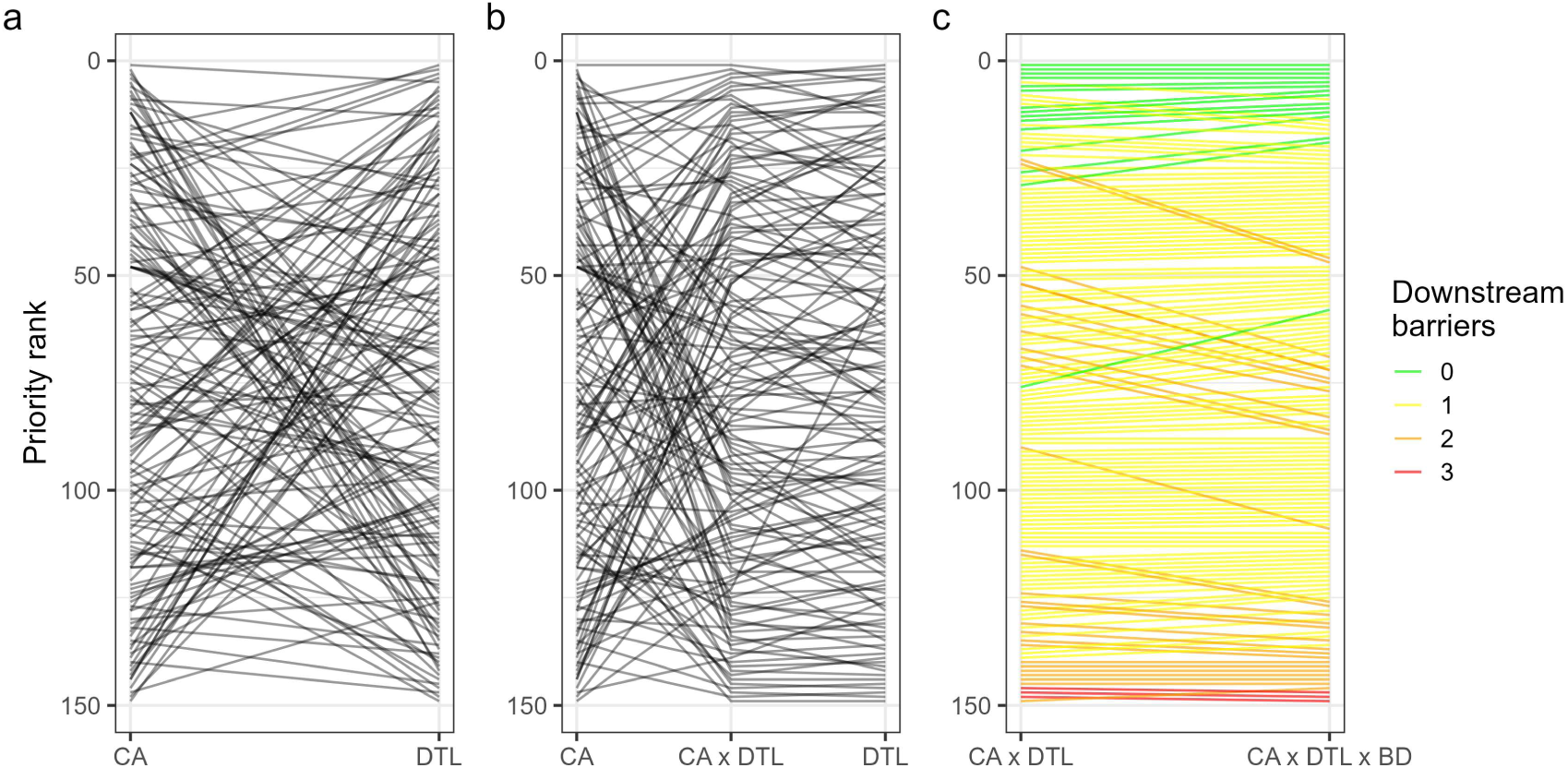
Rank changes of sites during geospatial bi-directional passage prioritisation, based on proxies for habitat quantity i.e. catchment area (CA), and downstream eel abundance, i.e. distance from tidal limit (DTL) separately (a), in combination (CAxDTL) (b), and with the number of barriers downstream (BD) introduced (c).

### Stage 2. Eel presence-informed directional passage prioritisation

Combining CAxDTLxBD scores and eel presence/absence resulted in significant re-ranking, with the largest rank change being 96 and eleven sites changing by 50 or more places, but there was still strong agreement between the two methods (Spearman’s rank, rho = 0.82, p < 0.001) (Figure 2). Indeed, pumping stations with small catchments and/or that were far from the tidal limit that contained eels were incorrectly ranked as low priority for downstream passage improvements using geospatial metrics alone. Conversely, sites without eels that were in large catchments and close to tidal water could have been incorrectly prioritised as high priority.

**Figure 2.**
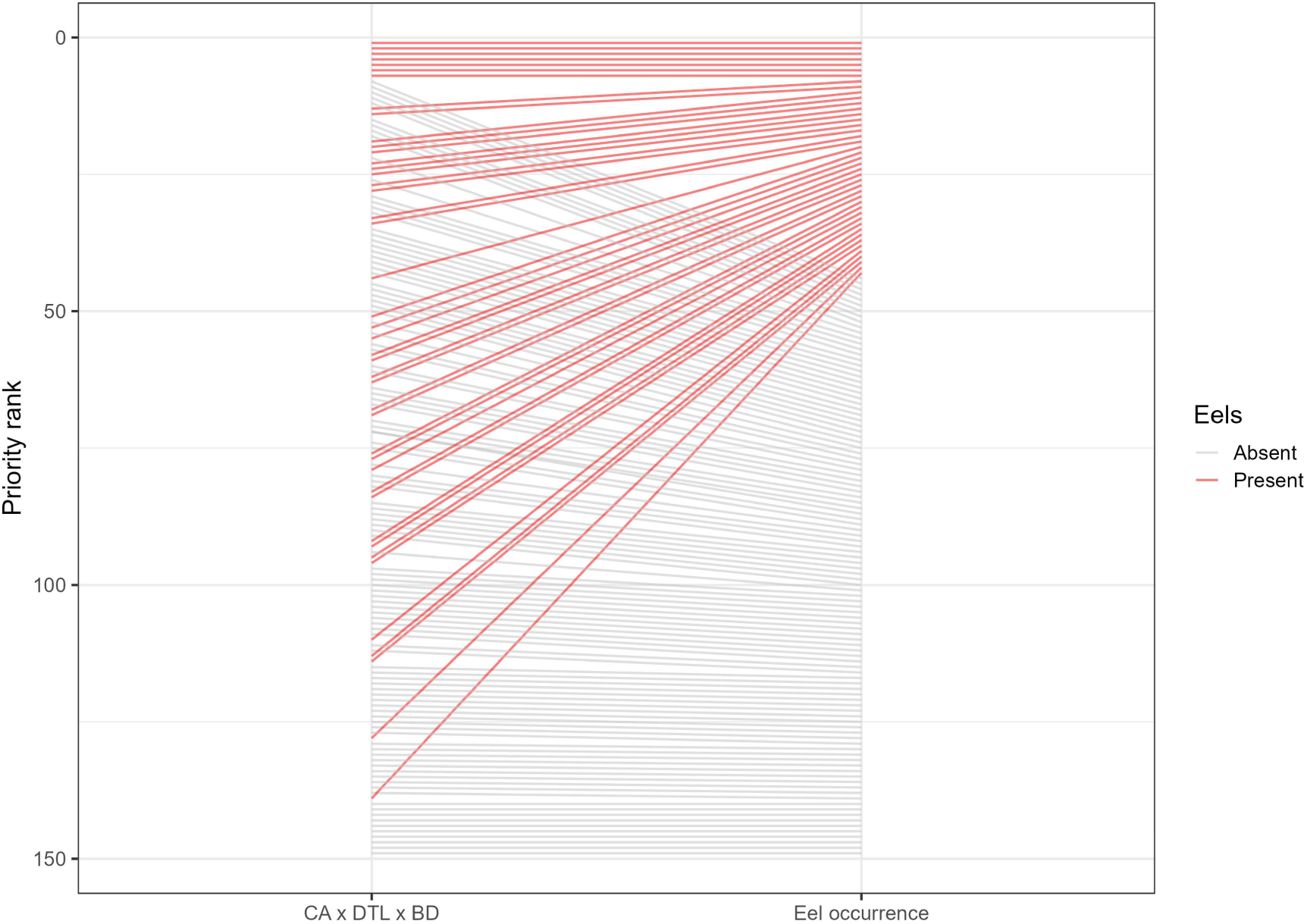
Rank changes of sites for bi-directional passage prioritisation based on the habitat quantity (CA), distance from the tidal limit (DTL), number of barriers downstream (BD) and the presence (red lines) and absence (grey lines) of eels.

### Stage 3. Prioritisation refinement using site-specific features

#### Downstream passage. Hydrological connectivity type (HCT); a proxy for upstream eel abundance

When incorporating connectivity as an eel abundance proxy into the formula (CAxDTLxBDxHCT), agreement remained strong (Spearman’s rank, rho = 0.97, p < 0.001), but significant re-ranking occurred (max. rank change of 25, four sites changed by 20 or more places, Figure 3).

**Figure 3.**
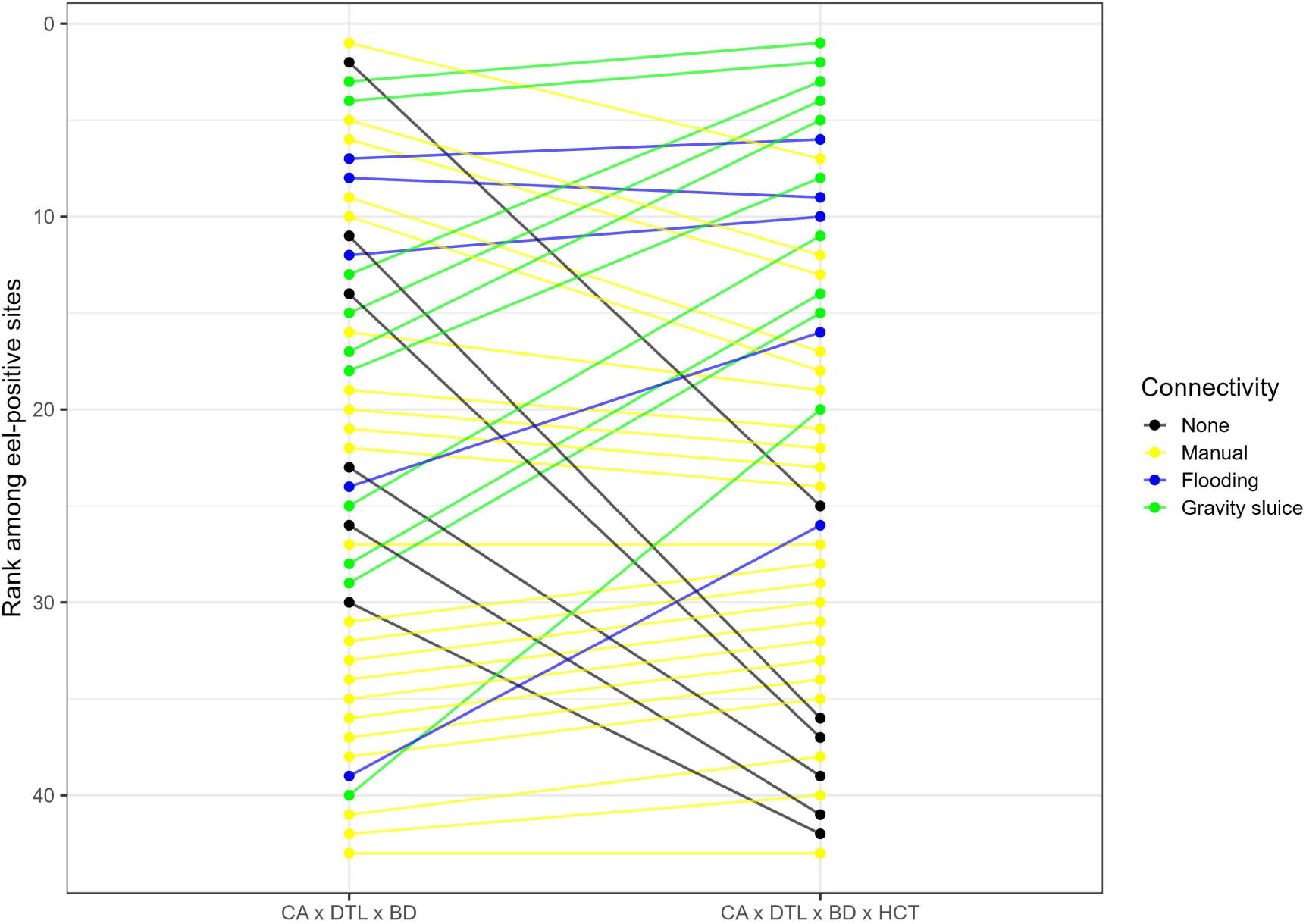
Rank changes of sites prioritised for downstream passage improvements where eels were detected based on the habitat quantity (CA), distance from the tidal limit (DTL), number of barriers downstream (BD) and connectivity as a proxy for eel abundance (HCT). Line colours denote connectivity types of sites.

#### Upstream passage. Winter fish richness (WFR); proxy for upstream habitat quality

At sites where eels were not detected, initial prioritisations for upstream eel passage based upon CA, DTL and BD were significantly altered by the incorporation of WFR (Spearman’s rank, rho = 0.50, p < 0.001) and significant re-ranking occurred, with fourteen sites changing by a rank of 50 or more (Figure 4).

**Figure 4.**
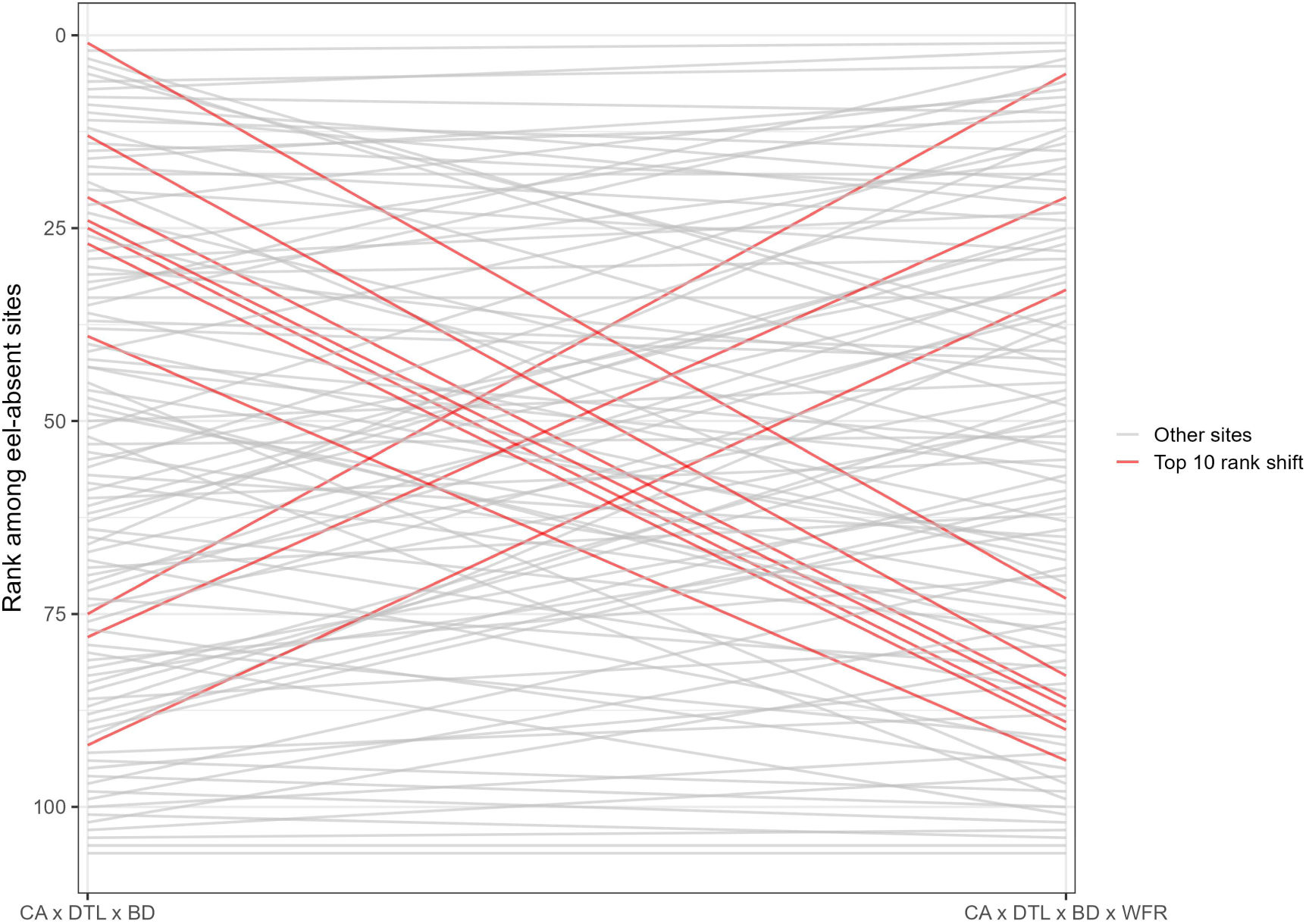
Rank changes of sites when prioritised for upstream passage improvements based on the habitat quantity (CA), distance from tidal limit (DTL), number of barriers downstream (BD) and winter fish richness (WFR) as a proxy for habitat quality. The ten largest rank shifts are marked in red.

### Final ranking

Several sites changed notably in downstream and upstream passage priority as additional factors were incorporated (Figure 5). For example, the pumping station ranked second based on geospatial metrics (CA, DTL and BD) and eel presence fell to 25^th^ for downstream passage given the absence of a known route for eels to enter the catchment. By contrast, the 113^th^ ranked site based on geospatial metrics increased to 40^th^ for downstream passage following eel detection and then to 20^th^ given the presence of a gravity sluice to provide a route for eel colonisation. For upstream passage, the site ranked 8^th^ based on geospatial metrics due to its large catchment and estuarine discharge was the top-ranked eel-absent site and fell to 73^rd^ when WFR was incorporated. Likewise, a site ranked 134^th^ increased to 92^nd^ and then 33^rd^ for upstream passage following the incorporation of eel-absence and WFR, respectively, while another site initially ranked 116^th^ increased to 75^th^ and then 5^th^. The final downstream and upstream rankings can be seen and are spatially visualised in Figure 5 and Figure 6, respectively.

**Figure 5.**
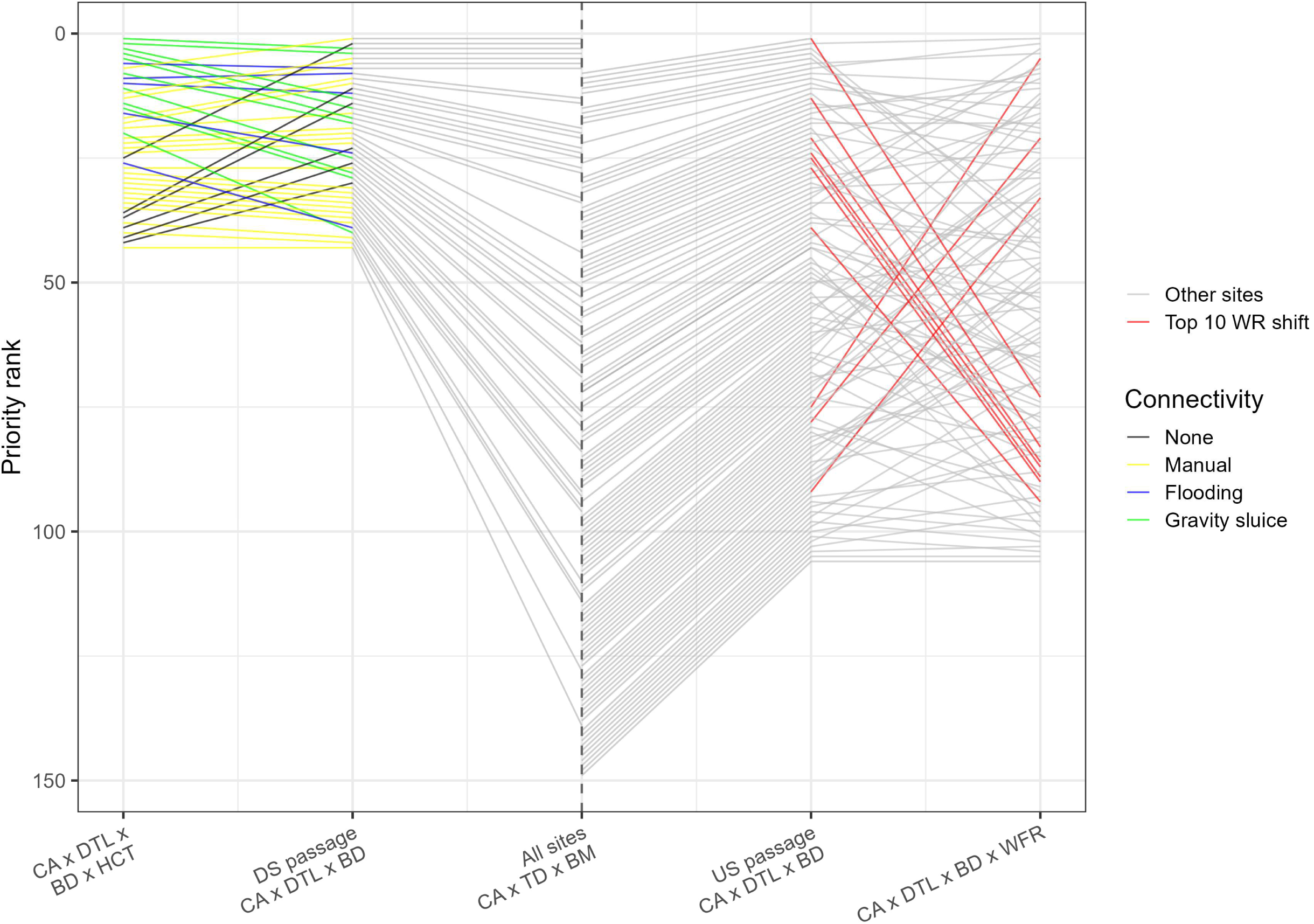
Rank changes of sites when prioritised for downstream and upstream passage (left and right of the dashed vertical line, respectively) based on the catchment area (CA), distance from tidal limit (DTL), number of barriers downstream (BD), hydrological connectivity (HCT) and winter fish richness (WFR).

**Figure 6.**
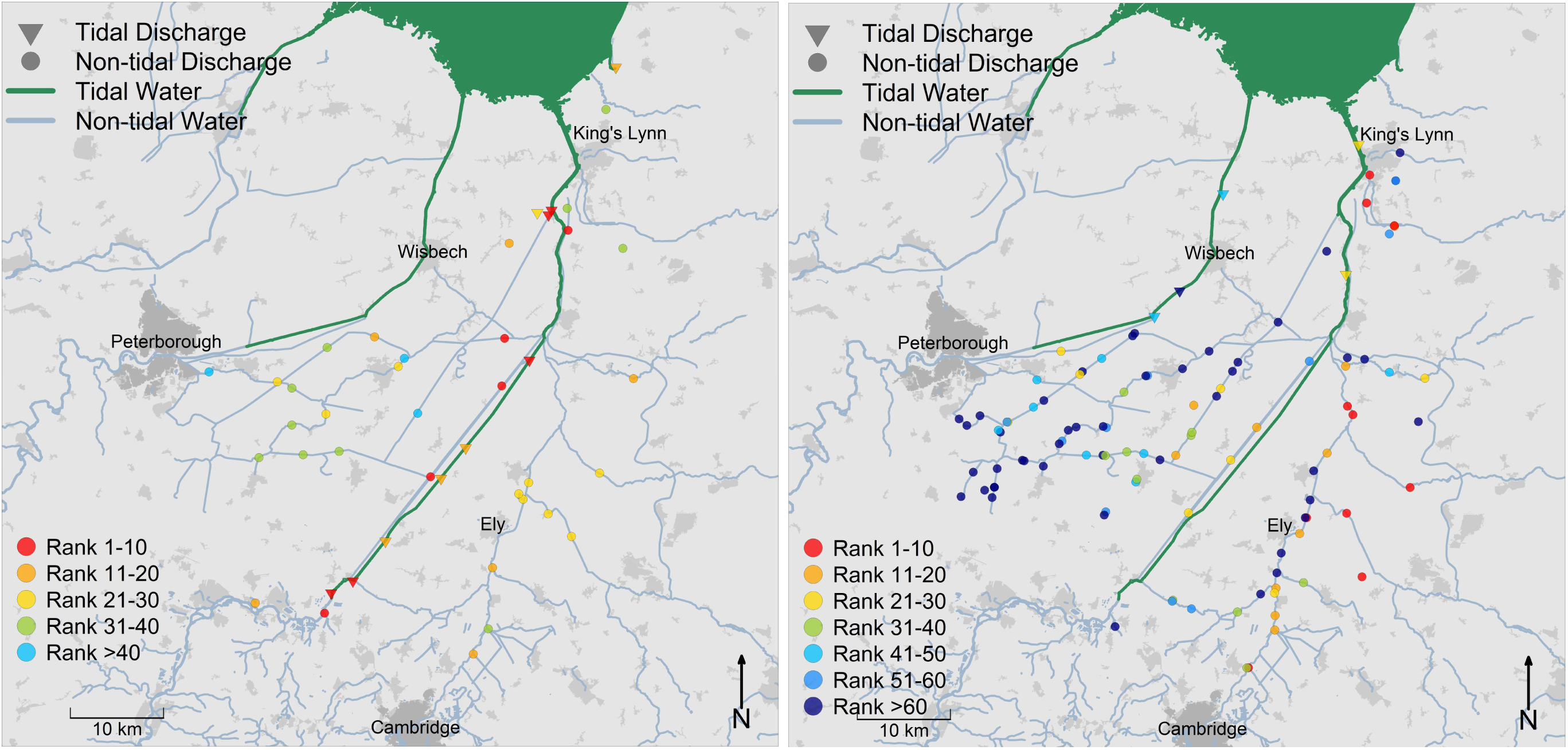
Spatial visualisation of final ranking for downstream (left) and upstream (right) eel passage prioritisation at pumping stations in The Fens.

## Discussion

Strategies to effectively manage and mitigate the impact of human activities on the natural environment are urgently needed to conserve biodiversity. To-date, efforts to protect Anguillid eels at hazardous infrastructure (e.g. pumping stations and hydropower) have relied on *ad hoc* site-specific assessments or desk-based prioritisations. Here, we uniquely incorporate geospatial, present-day biological data and site-specific features to iteratively prioritise critically endangered European eel protection at pumping stations in a former wetland at ecosystem-scale. The value of understanding and incorporating all levels of data was evident, often significantly altering rankings. Incorporating eel occurrence also uniquely enabled downstream and upstream passage to be prioritised separately to allow for bespoke conservation outcomes.

Previous fish passage prioritisations range between using simple geospatial metrics (Kocovsky et al., 2009; Solomon & Wright, 2012), incorporating historic fish community data (Clavero & Hermoso, 2015; King et al., 2017) and more detailed site-specific assessment with contemporary fish community, passability and habitat data (Nunn & Cowx, 2012). However, to our knowledge, previous fish passage prioritisations have not quantified how site priorities change as successive sources of biological and site-specific information are incorporated. Similarly to Kujala et al. (2018), which focused on large-scale terrestrial conservation areas, we found that incorporating certain additional factors into prioritisations had profound effects. Most notably, incorporating present-day knowledge of eel occurrence significantly influenced the prioritisation based on geospatial factors. Indeed, sites close to the tidal limit and large catchment area were often eel-less and, conversely, eels were detected in small sites far from tidal water. This highlights the complex nature of species distributions in fragmented and hydrologically complex systems (Wilson et al., 2016; Griffiths et al., 2025a). Thereafter, significant re-ranking of downstream passage occurred according to the type of hydrological connectivity, as a proxy for eel abundance, and upstream passage occurred according to fish species richness, as a proxy for habitat quality.

A hierarchical approach of ‘remove’ the pumping station, ‘replace’ harmful infrastructure with fish-friendly equivalents/alternatives or ‘mitigate’ the effects by modifying infrastructure and/or operation should be applied to pumping stations to improve two-way connectivity (Bolland & Franklin, 2025). In this study region, The Fens, socio-economic drivers largely prevent pumping stations from being removed and considerations for where, how and when to replace or mitigate to remediate downstream and upstream passage are inherently different. Hereafter, we consider the subtleties associated with each and practicalities of applying the findings of this prioritisation in the real-world.

### Downstream passage

European eels can be directly (e.g. blade strike as they pass through traditional pumps) or indirectly (e.g. predation during passage delays) injured or killed as they move seawards to complete their catadromous lifecycle (Bolland et al., 2019). Providing safe and timely downstream passage at pumping stations is urgently required (Evans et al., 2024). In the absence of present-day eel distribution data, it must be assumed eels are present upstream of all pumping stations, as per their historic distribution. However, here, contemporary eDNA data was used to inform this prioritisation; European eel were only present upstream of 28.3% of pumping stations (Monaghan et al., 2026). It is hence recommended that, in the short-term, eel protection should be focused on pumped catchments that contain eels rather those that do not.

Replacing traditional pumps and other harmful infrastructure with fish-friendly equivalents/alternatives is presently considered the preferred eel downstream passage solution. However, further standardised research is urgently needed to quantitatively demonstrate that eels pass through stations with fish-friendly pumps without migration delay, injury or mortality (Evans et al., 2024). Furthermore, it may take many years for pumps to be replaced. Alternative approaches to mitigate downstream passage should be sought in the interim. Temporary screens or nets could be installed to prevent eel entrainment, provided flood-relief remains unhindered and eels do not become impinged (Carter et al., 2023a). Many (10 of 43; 23.3%) of the sites where eels were detected have a gravity sluice, and thus operations could be manipulated to quickly provide a cost-effective alternative downstream passage route (Carter et al., 2023a, 2025). At sites without a gravity sluice, a bespoke bypass could be provided, but operation is inherently challenging at pumping stations given the requirement to move water to a higher altitude. Crucially, the findings from this prioritisation will enable more targeted conversations with site owners about the feasibility of interim and more permanent eel protection measures.

In theory, conservation efforts should focus on the largest populations of rare or endangered animals, and thus downstream passage should be prioritised in catchments with the largest eel stocks. The eDNA data used here (Monaghan et al., 2026) revealed eel presence/absence with a high level of sensitivity relative to traditional fish sampling methods (Griffiths et al., 2020) and confidence (Griffiths et al., 2025b), but eel density across catchments remains unknown. Consequently, the type of hydrological connectivity was used as a proxy for eel abundance, informed by (Monaghan et al., 2026). Doing so allowed sites with similar geospatial characteristics to be assessed on a more site-specific basis. Indeed, four sites changed ranking by at least twenty places. Notwithstanding, capture-based fish surveys in eel-positive catchments could be employed to compare size, structure (e.g. lengths, weights and life-stage) and condition of populations (Degerman et al., 2019) and further inform decision-making.

### Upstream passage

Remediating upstream passage into locations that once supported eel is generally seen as an appropriate conservation action (Clavero & Hermoso, 2015). It mirrors many large-scale conservation initiatives, such as the Water Framework Directive, which emphasise the importance of maximising access to habitat via improving fish passage at migration barriers (Belletti et al., 2020). However, in the case of pumping stations, practitioners must proceed with caution. Firstly, a safe, highly efficient and timely downstream passage solution must be implemented for eels that subsequently colonise habitats upstream of the pumping station; the ranking for remediating upstream passage is hence also a ranking for remediating downstream passage. Secondly, the cause of eel absence must also be established. Eel absence could be attributed to pumping stations (and associated flood banks) being a complete barrier to upstream migrating eels, and thus remediating upstream passage may lead to increased escapement. Alternatively, eel absence may be attributed to degraded upstream habitat quality, such as low water levels and low dissolved oxygen during eutrophic events (Dawson, 2002), and thus remediating upstream passage may unintentionally kill eels, potentially exacerbating eel decline. Hence it is necessary to consider the long-term implications of re-introducing a critically endangered species into habitat that may be sub-optimal and vulnerable to climate change and other anthropogenic pressures (van Puijenbroek et al., 2021).

During this study, upstream eel passage was prioritised at larger catchments closer to the tidal limit, provided their habitat can support a diverse fish community (i.e. high winter fish richness). The largest catchments without eels offer the greatest potential to contribute to stock recovery and may be more resilient to intermittent or systemic water quality issues (Beevers et al., 2021). Sites closer to the tidal limit were prioritised given greater numbers of juveniles present in tidal waters and a strong propensity to migrate upstream (Bevacqua et al., 2011). Eels have a broad range of environmental tolerances (Nälsund et al., 2022), but it is recommended upstream passage improvements should nonetheless be prioritised into areas with high-quality habitats due to its potential positive effect on eel survival and condition, ultimately increasing the likelihood of reproductive success (Belpaire et al., 2011). It was not feasible due to time, budget and practical constraints to quantify site-specific physical habitat or water quality. Instead, winter fish richness (as a conservative metric for fish community persistence following temporary hydrological connectivity; Monaghan et al., 2026) was used as a biotic indicator of habitat quality (Karr, 1981; Pont et al., 2006). The changes in upstream passage rankings following its inclusion demonstrate that sites with similar geographical characteristics can have considerably different habitat quality and thereby improved the prioritisation.

Upstream passage into pumped catchments is not an obligatory requirement for juvenile catadromous species to complete their lifecycle, which is unlike upstream migrating adult anadromous species attempting to reach spawning grounds (Verhelst et al., 2021). Indeed, European eel can live and mature in coastal waters, estuaries, rivers, wetlands and lakes, although the relative contribution each habitat makes to the adult stock remains poorly understood (Righton et al., 2016, 2021). Consequently, it is recommended that upstream passage should not be provided into pumped catchments with the poorest quality habitat, unless the underlying mechanisms for habitat deterioration can be reversed (Alp & Melching, 2011). However, there is a risk that not providing upstream passage into catchments with the poorest habitat promulgates a disincentive for pumping station owners to rehabilitate the habitat.

The mechanism to remediate upstream passage at pumping stations, if at all, depends on the type of hydrological connectivity. Elver passes (Solomon et al., 2004) could be employed at pumping stations with no hydrological connectivity or that are rarely connected when the downstream river floods. Conversely, stations with operational gravity sluices could be excluded from upstream passage remediation, given a viable upstream passage route already exists (Piper et al., 2012; Monaghan et al., 2026). Indeed, eel absence may reflect the location in the upper catchment and thus low number of eels in the downstream waterbody with low propensity to migrate upstream (Yokouchi et al., 2022). As such, if eel recruitment improves in the future, these sites may be colonised by eels without the assistance of elver passes. Sites with mechanisms to manually transfer water upstream, such as for irrigation purposes, have to potential to provide upstream eel passage provided they are operated at times when upstream migrating eels are known to migrate. Conversely, lowly ranking sites for upstream passage with poor habitat and existing gravity sluice or manually transfer mechanism may need to be screened (Carter et al., 2023a) to prevent eels entering a catchment they are unlikely to emigrate from.

At sites where eels are present, it is recommended that upstream passage is remediated at the same time as downstream passage. However, this may not be necessary at the top ranked sites for downstream passage as they have a gravity sluice, and thus a reliable immigration route already exists. Notwithstanding, sites that contain eels may have poor quality upstream habitat and therefore the ranking for downstream passage may not necessarily agree with priorities for upstream passage. Therefore, as with providing upstream passage into catchments without eels, practitioners must proceed with caution to avoid remediating upstream passage into habitat that may be sub-optimal and may unintentionally kill eels.

### Real-world application

This investigation exclusively considered ecological factors to prioritise eel protection at pumping stations and thus does not reflect the realities of funding mechanisms nor practicalities of managing pumping stations, such as site-specific or drainage board refurbishment schedules. The challenge of funding fish protection in the real-world has also been reported for other types of infrastructure due to conflicting priorities when considering ecological benefit, costs and works schedules independently, rather than together (Kemp & O’hanley, 2010; Garcia de Leaniz & O’Hanley, 2022). Indeed, given the significant cost and engineering challenge associated with retrofitting eel protection at existing sites, upstream and downstream eel passage is currently implemented according to Environment Agency and Drainage Board programmes of non-eel-related works. Moving forwards, this prioritisation can be used to support decision-making, especially at sites not yet included in a programme of works. For example, we recommend high-priority downstream passage sites that are not due for refurbishment in the near-future, potentially decades, should be considered sooner than sites without eels to prevent large numbers of eels being at risk of entrainment for many years (Bolland et al., 2019; Evans et al., 2024). Conversely, lowly ranking sites for upstream passage remediation due to poor habitat quality should be considered later, potentially using the time in the interim to improve the habitat. The prioritisation can also be used to inform where site-specific physical habitat and water quality assessments could be performed to better understand the underlying reasons for eel absence and further inform upstream passage decision making. In doing so, the results of this prioritisation will deliver the greatest possible eel conservation benefits within the current regulatory framework, delivery schedules and funding landscape.

## Conclusion

The socioeconomic reliance on pumping stations, coupled with a limited present-day distribution of critically endangered European eel and the technical, logistical and financial challenges associated with remediating connectivity necessitates strategic allocation of limited resources to conserve eels and comply with legislation (EC Regulation No. 1100/2007). The reproducible and transparent prioritisation produced here integrated geospatial, contemporary biological data and site-specific features to ensure safe downstream passage is prioritised for eels currently inhabiting pumped catchments and juvenile eels colonise the highest quality, most resilient habitats to maximise future escapement. Indeed, incorporating eel occurrence and wider fish community significantly altered site rankings compared with approaches based solely on catchment area and distance from tidal water, demonstrating that map-based prioritisation alone can overlook sites with the greatest conservation potential. For upstream passage, the cause of eel absence must be established to ensure eels are not provided access into degraded habitat quality where they are unable to persist during periods of hydrological disconnection. The findings from the prioritisation can be integrated with ongoing programmes of infrastructure refurbishment to deliver the greatest eel conservation benefits according to current funding mechanisms and real-world practicalities. The approach is also transferable to other heavily modified landscapes supporting anguillid eels globally.

## Acknowledgements

We thank the drainage boards for their cooperation. We thank the Environment Agency and the University of Hull for funding the study.

## Author Contributions

N.P.G, A.D.N, R.M.W and J.D.B developed the study. R.M.W and J.D.B acquired funding. A.I.T.M and G.S.S conducted data analysis. A.I.T.M drafted the first manuscript. All authors contributed to manuscript revisions.

## Data Availability

Data and scripts have been made available on Zenodo (https://doi.org/10.5281/zenodo.24118514).

## Use of AI

ChatGPT (OpenAI) was used for assistance with coding for data analysis and language editing for clarity. Outputs were verified by authors.

## Notes

### Competing Interest Statement

The authors have declared no competing interest.

